# Young chicks quickly lose their spontaneous preference to aggregate with females

**DOI:** 10.1101/2020.08.28.272146

**Authors:** Virginia Pallante, Daniele Rucco, Elisabetta Versace

## Abstract

It is not clear when and how animals start to discriminate between male and female conspecifics and how this distinction drives their social behaviour. A recent study on pheasants found that one-week-old chicks (*Phasianus colchicus*) preferentially aggregated with same-sex peers and this trend became more pronounced through development, suggesting that sexual segregation increases during ontogeny. However, it remains unclear whether this ability depends on experience or develops spontaneously. Using a similar experimental protocol, we investigated whether sex discrimination is present at birth in domestic chickens (*Gallus gallus*) by testing the aggregation preferences of young chicks with clutch mates. We measured the amount of time spent close to male and female conspecifics in visually inexperienced chicks. Soon after hatching, both males and females preferentially aggregated with females. To clarify whether the experience with conspecifics modifies the initial preference for females we used an imprinting procedure. We exposed chicks to conspecifics of the same sex, different sex or both sexes for three days and then tested their preferences to aggregate with males or females. No sex preference was observed after three days of imprinting exposure. The disappearance of the initial sex preference shows that, although chicks can discriminate between conspecifics of different sex, sex segregation does not influence aggregation in the first week of life. We suggest that the absence of sexual assortment in the first week of age can enhance the social cohesion of the flock.

## Introduction

Predispositions that support social life orienting the behaviour to social partners have been widely documented in precocial animals, namely species that have a mature sensory system and move autonomously soon after hatching or birth (Versace et al. 2017a). For instance, in the first hours after hatching, domestic chicks preferentially approach biological motion (Vallortigara et al. 2005; Vallortigara and Regolin 2006) and motion dynamics associated with animate objects such as changes in speed (Rosa-Salva et al. 2016; Versace et al. 2019), quails are attracted by engaging social partners (ten Cate 1986) and, similarly to chicks and ducklings, preferentially approach their own species maternal call (Gottlieb 1965, 1974; Park and Balaban 1991). Chicks are attracted to face-like stimuli and cues present in the head region (Johnson and Horn, 1988; Rosa-Salva et al. 2010, 2019; Versace et al. 2017a) or specific colour portions of the spectrum, such as the red and the blue regions (Hess and Gogel 1954; Hess 1956; Miura et al. 2020). On the one hand, this evidence shows abilities to discriminate between subtle features of the stimuli. On the other hand, it shows the existence of preferences that help young animals to orient towards cues that signal the presence of animate, living beings, that are not species-specific (reviewed in Di Giorgio et al. 2017 and Rosa-Salva et al. under review). Much less is known on preferences that young animals might have within their own species, such as the preference to aggregate with males and females. In Galliformes, sexual segregation is apparent when animals are sexually dimorphic (Whiteside et al. 2018), so that when sexual dimorphism becomes clear animals preferentially aggregate with individuals of the same sex. However, little is known on when the ability to differentiate between males and females emerges, and whether in the first days of life chicks preferentially aggregate with one sex or the other. Two main scenarios are possible: from the early stages of life, chicks prefer to aggregate with same-sex conspecifics, or the preference for one of the two sexes may be acquired later through experience.

A study on early behavioural differences between sexes has been conducted in pheasants (*Phasianus colchicus*) (Whiteside et al. 2017). Authors investigated sex preference in young chicks to understand the proximate mechanisms driving sexual segregation. They found ontogenetic changes in the early propensity of male and female chicks to aggregate with different sexes. In the first week after hatching, before sexual dimorphism becomes marked, females preferred approaching their own sex. This preference increased during ontogeny in both male and female chicks (Whiteside et al. 2017). A further study showed that 10-week-old chicks released into the wild exhibit patterns of sexual segregation in their social preference at feeding sites. Such preference becomes more pronounced after the first month from the release (Whiteside et al. 2019). However, it remains to be shown whether the preference to aggregate with one sex or the other is present at birth or acquired later through experience with conspecifics of different sex.

To understand whether sex discrimination is a spontaneous ability that does not require experience, we used a procedure similar to that used by Whiteside et al. (2017) to test the aggregation preferences of young chicks (*Gallus gallus*) in an aggregation test, soon after hatching (Experiment 1). Because we observed a spontaneous preference of newly hatched chicks to aggregate with females, in line with the preference previously observed in one-week old pheasants (Whiteside et al. 2017), we used a filial imprinting paradigm to investigate the influence of experience with members of the two sexes.

Filial imprinting is a fast learning process that takes place in the early stages of life and enables naïve individuals to learn the features of their social partners through mere exposure, driving attachment and preferences (Bateson 1966; McCabe 2013; Vallortigara and Versace 2018). During the imprinting phase, individuals acquire and memorize the particular features of the imprinting stimulus and build a representation that supports a subsequent preference to aggregate with it (Bateson 1966, 1990; Bolhuis 1991; Bischof 2003; Matsushima et al. 2003; Horn 2004). An early exposure to parents and peers also affects sex preferences once sexual maturity has been reached (Bateson 1966; Clayton 1989). Indeed, learning the morphological features of the conspecifics during the juvenile period will drive sex discrimination and preference towards same species individuals later in life (Sherrod 1974; Worseley 1974; Cheng et al. 1979). The exposure to different sexes in the imprinting phase may also affect sex preference at maturity. For instance, in chickens and pheasants the lack of experience with opposite sex in the first stages of life seems to influence opposite sex preference and mating success in adulthood (Leonard et al.1993; Madden and Whiteside 2013). In addition, male and female domestic chicks differ in their preference towards the imprinting stimuli (Vallortigara and Andrew 1991; Vallortigara 1992; Versace et al. 2017b). For example, males and females differ in their behaviour towards familiar and unfamiliar stimuli in terms of latency and time spent near the stimulus (Vallortigara 1992) and overall attraction, with a male preference towards unfamiliar stimuli (Versace et al. 2017b). This latter result is consistent with the so called “slight novelty effect” (Jackson and Bateson 1974; Bateson 1979; Versace et al. 2006), indicating a preference for a novel stimulus compared to a familiar one and observed particularly in males (Vallortigara and Andrew 1994). The differences between sexes and across species (see *Coturnix japonica,* Launay et al. 1993) in the social response may rely in the species’ social system that shaped gregariousness predisposition in different sexes (Vallortigara et al. 1990).

Do chicks maintain or modify their spontaneous preference for females due to their exposure to same/difference sex conspecifics? To investigate the effect of social exposure on sex aggregation, we manipulated chicks’ experience by imprinting them with chicks of the same sex (Experiment 2), different sex (Experiment 3) or mixed sexes (Experiment 4) and tested their preference to approach male or female chicks. Finally, confirmed the absence of the sexual dimorphism between male and female chicks using physical traits (weight) and the level of activity (distance moved).

## Materials and Methods

### Subjects and rearing conditions

In four experiments, we tested 872 chicks of domestic fowl (*Gallus gallus*) of the strain Ross 308 (see Table 1). This strain is characterized by a sexual dimorphism at birth in the disposition of the primary covert feathers. The eggs were obtained from a commercial hatchery (Agricola Berica, Montegalda, Italy) and incubated in darkness at 37.7 °C and 40% of humidity for the first 18 days and 60% for the remaining 3 days before hatching.

**Table 1.**
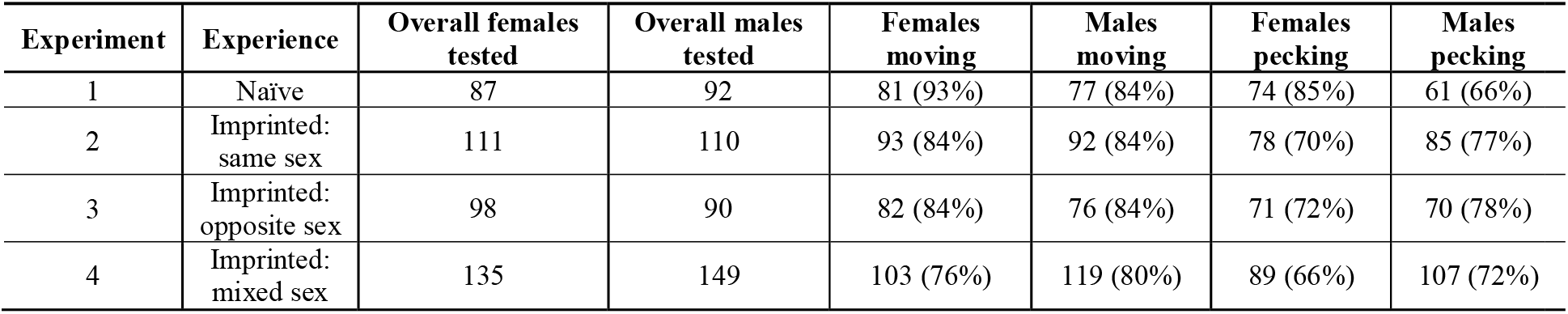
Chicks used in each experiment. For each experiment, the table shows how many males and females have been tested, how many males and females made a choice (moving chicks) and the percentage with respect to the whole number of tested chicks, how many chicks made a choice and interacted with the conspecifics and the percentage with respect to the whole number of tested chicks

| Experiment | Experience | Overall females tested | Overall males tested | Females moving | Males moving | Females pecking | Males pecking |
| --- | --- | --- | --- | --- | --- | --- | --- |
| 1 | Naïve | 87 | 92 | 81 (93%) | 77 (84%) | 74 (85%) | 61 (66%) |
| 2 | Imprinted: same sex | 111 | 110 | 93 (84%) | 92 (84%) | 78 (70%) | 85 (77%) |
| 3 | Imprinted: opposite sex | 98 | 90 | 82 (84%) | 76 (84%) | 71 (72%) | 70 (78%) |
| 4 | Imprinted: mixed sex | 135 | 149 | 103 (76%) | 119 (80%) | 89 (66%) | 107 (72%) |

In Experiment 1, eggs were hatched either in individual plastic boxes (11 cm x 8.5 cm x 14 cm) or in group physical contact. Since no difference was observed between the two conditions (see Supplementary material), in subsequent experiments chicks were hatched in groups. The same hatching conditions were used for Experiment 2, 3 and 4.

We analysed the performance of chicks that made a choice (active chicks that moved out of the staring zone, see Experimental procedure) to investigate their preference for male and female chicks. Overall, we analysed the behaviour of 635 chicks, 323 males and 312 females (Table 1).

Among active chicks, we further analysed those that interacted with the stimuli touching or pecking the plastic mesh that delimited the area with the stimuli. Each chick was tested once. After the test, chicks were housed in groups in cages (125.5 cm x 76 cm x 45 cm), with water and food available ad libitum, in a 14:10 light: dark regime at 28-30 °C and donated to local farmers.

### Test apparatus

The test apparatus was a white corridor (100 cm x 35 cm x 31 cm) divided in three different chambers: a central chamber (60 x 35 cm) and two lateral chambers (20 x 35 cm each), see Fig. 1.

**Figure 1.**
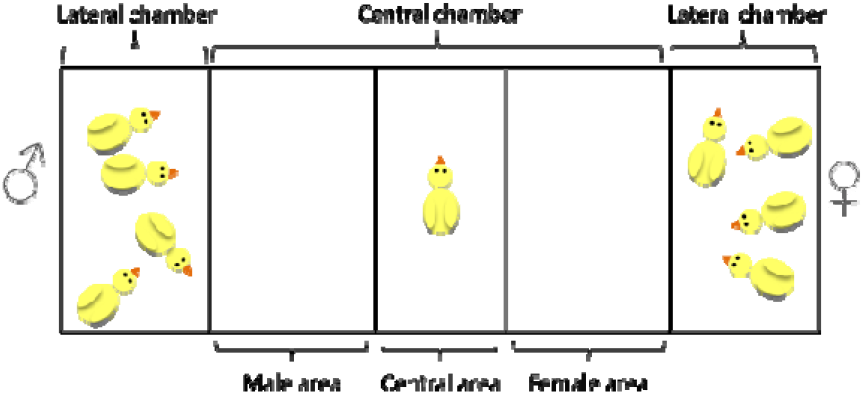
Illustration of the test apparatus. The right/left position of male and female chicks used as stimuli was counterbalanced between subjects.

In the two lateral chambers we placed the experimental stimuli: four female chicks or four male chicks from the same hatch of the tested chicks. The right/left position of males and females was counterbalanced between trials. The three chambers were separated by a transparent acrylic mesh. The central chamber was additionally divided into three areas: left area (male area in the figure, 21.5 x 35 cm), central area (17 x 35 cm) and right area (female area in the figure, 21.5 x 35 cm). These areas were indicated by a line on the floor. A video-camera located above the apparatus recorded the test. Two stripes of LED were placed on the top of the two lateral chambers to illuminate the apparatus (BB-ER1020W0, SMD 5050, 1080 lm/m).

### Experimental procedure

#### Experiment 1: Naïve chicks

In Experiment 1, chicks’ preferences were tested soon after birth. Chicks were individually transported from the incubator to the experimental room inside a dark plastic box, sexed and tested. At the beginning of the testing phase the chick was placed facing the long side of the apparatus, on the opposite side of the experimenter. We recorded the time the chick spent in each area of the central chamber for six consecutive minutes. Each chick was scored in one area when the whole body entered that area. We measured the preference for females as

The positive values indicate a preference towards females, the negative values a preference for males.

#### Experiments 2-4: Imprinting experiments

In Experiments 2-4, chicks were imprinted for 3 days and then tested on the fourth day of life. Newly-hatched chicks were taken from the incubator and individually transported into the adjacent maintenance room, sexed and transferred in a group of five animals into an imprinting cage (28 cm x 40 cm x 32 cm). Chicks were imprinted for three days with same/opposite-sex chicks of the same hatch: in Experiment 2 with four same-sex chicks, in Experiment 3 with four opposite-sex chicks, in Experiment 4 with two opposite-sex and two same-sex chicks. In the imprinting cages, water and food were available ad libitum, the temperature was constantly at 28-30°C and the light:dark conditions were 14:10 hours. Chicks could hear but not see chicks placed in the other cages. The chicks used as stimuli were reared in the same conditions of the testing chicks.

### Analysis of distance moved and body weight

For each condition we analysed the distance moved by the focal subject (in centimetres) for ten videos for each sex. Using the software Ethovision (Noldus), we tracked the distance moved by the testing chicks during the testing phase. At the end of the experiment we measured the weight of a sample of chicks using an electronic scale: 79 females and 58 males in Experiment 1 and 277 females and 280 males in Experiment 2, 3 and 4.

### Statistical analysis

We analysed the preferences for females using an ANOVA with Sex as between-subjects variable and Time (minutes 0-6) as within-subjects variable. We ran two analyses: one on all chicks that approached the lateral choice sectors (active chicks) and one on chicks that touched the transparent divider and hence showed greater motivation to approach the social partners (see Table 1). We run an ANOVA to test the effect of the Time (minutes 0-6) as within-subjects variable and the

Sex of the testing chick as between-subjects variable on preference for females. We evaluated the significance of the preference for females against the chance level with one-sample t-tests. We analysed the distance using an ANOVA with Time (minutes 0-6) as within subjects-variable and Sex as between-subjects variable. We compared the distance moved in each experiment running an ANOVA with Experiment as between-subjects variable and used the Tukey’s test for post-hoc comparisons. We compared the weight of males and females with Welch two sample t-tests. All the analyses where performed with R (version 3.4.4), with alpha level set to 0.05.

## Results

### Experiment 1. Naïve chicks

When analysing all active chicks, we found no significant main effect of Sex (F_1, 156_ = 1.851; p = 0.176) and no significant interaction Sex x Time (F_5,780_ = 0.347; p = 0.884). However, we found a significant effect of Time (F_5,780_ = 2.286; p = 0.044), with chicks increasingly spending more time with females during the test, particularly in the fourth minute compared to the first minute (t_199.03_ = −2.954; p = 0.003, see Supplementary Fig. 1). When we restricted our analysis to highly motivated chicks, we observed no significant main effect of Sex (F_1,132_ = 0.766; p = 0.383) or Time (F_5,660_ = 2.154; p = 0.057) and no significant interaction Sex x Time (F_5,660_ = 0.233; p = 0.948). Hence, we analysed the overall preference and observed that newly hatched chicks prefer to spend more time with females (t_133_ = 3.204; p = 0.002, Fig. 2a).

**Figure 2.**
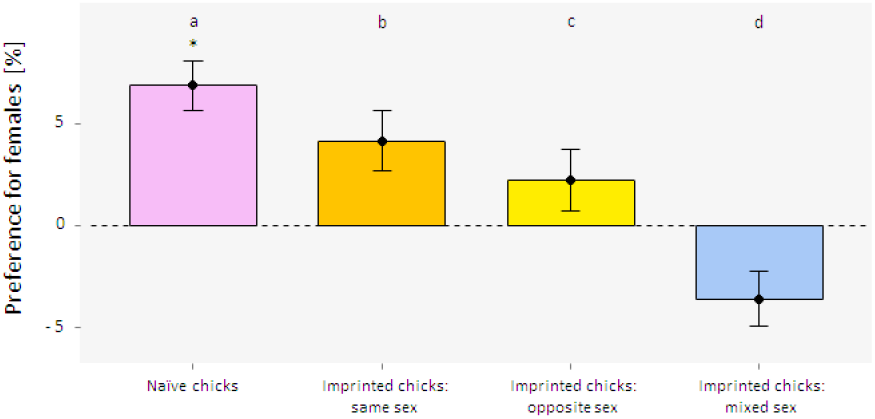
Barplots showing the overall preference for females in highly motivated chicks (mean ± SEM). **a.** Naïve chicks: Experiment 1; **b.** Imprinted chicks: same sex: Experiment 2; **c.** Imprinted chicks: opposite sex: Experiment 3; ; **d.** Imprinted chicks: mixed sex: Experiment 4

### Experiment 2. Chicks imprinted with same-sex conspecifics

When we analysed the behaviour of active chicks, we observed no significant effect of Sex (F_1,181_ = 0.087; p = 0.769) and Time (F_5,917_ = 0.784; p = 0.561). We found a significant interaction Sex x Time (F_5,917_ = 2.565; p = 0.026), with males significantly prefer to aggregate with females in the first minute (t_181.82_ = −2.023, p = 0.044; see Supplementary Fig. 2). No significant effect of Sex (F_1,159_=0.009; p = 0.925) or Time (F_5,807_ = 0.818; p = 0.537) and no significant interaction Sex x Time (F_5,807_=1.508; p =0.185) were found in highly motivated chicks. We did not find any overall preference (t_162_ = 1.416, p = 0.159; Fig. 2b).

### Experiment 3. Chicks imprinted with opposite-sex conspecifics

We found no effect of Time (F_5,790_ = 1.632; p = 0.149), Sex (F_1,151_ = 0.732; p = 0.394) and the interaction Sex x Time (F_5,790_ = 0.767; p = 0.573) on the preference for females of active chicks. Chicks did not show an overall preference to aggregate with females (t_158_ = 1.041, p = 0.299). When we considered highly motivated chicks only, no effect of Time (F_5,699_ = 1.362; p = 0.237), Sex (F_1,134_ = 1.104; p = 0.295) and the interaction Sex x Time (F_5,699_ = 0.852; p = 0.513) were observed in the preference for females. Chicks did not show an overall preference to aggregate with females (t_140_ = 0.740, p = 0.460; Fig. 2c).

### Experiment 4. Chicks imprinted with mixed-sex conspecifics

We did not find any significant effect for Sex (F_1,219_ = 0.300; p = 0.584), Time (F_5,1095_ = 1.337; p = 0.246) and Sex x Time (F_5,1095_ = 1.135; p =0.340) in active chicks. We did not find an overall preference for females (t_220_ = −1.537, p = 0.126). Similarly, we found no significant effect for Sex (F_1,194_ = 0.578; p = 0.448), Time (F_5,970_ = 1.004; p = 0.414) and Sex x Time (F_5,970_ = 1.356; p = 0.238) in highly motivated chicks. No overall preference for females was observed (t_195_ = −1.356, p = 0.177; Fig. 2d).

The preference for females differed across experiments (Fig. 2). When comparing different experiments across time and between sexes in active chicks, we found a significant effect of Experiment (F_3,2400_ = 18.455; p<0.001), Sex x Time (F_5,1540_ = 2.709; p = 0.019) and the interaction Experiment x Sex x Time (F_15,1540_ = 2. 84; p = 0.002). We found an effect of the Experiment in the preference to aggregate with females for active highly motivated chicks (F_3,1920_ = 10.619; p<0.001).

### Distance moved

In Experiment 1 we found a significant effect of Time (F_118_ = 38.607; p < 0.001) but no effect of Sex (F_1,18_ = 0.341; p = 0.567) or of the interaction Sex x Time (F_1,18_ = 0.319; p = 0.579) (Fig. 3). In Experiment 2 we did not find any significant effect of Time (F_1,18_ = 1.019; p = 0.326), Sex (F_1,18_ = 1.421; p = 0.249) or of the interaction Sex x Time (F_1,18_ = 0.032; p = 0.861) (Fig. 3). In Experiment 3 Time had a significant effect (F_1,18_ = 14.809; p = 0.001), but not Sex (F_1,18_ = 0.519; p = 0.481) or the interaction Sex x Time (F_1,18_ = 0.209; p = 0.653) (Fig. 3). In Experiment 4 we did not find any effect of Time (F_1,18_ = 0.452; p = 0.510), Sex (F_1,18_ = 1.759; p = 0.201) or interaction Sex x Time (F_1,18_ = 1.157; p = 0.296) (Fig. 3).

**Figure 3.**
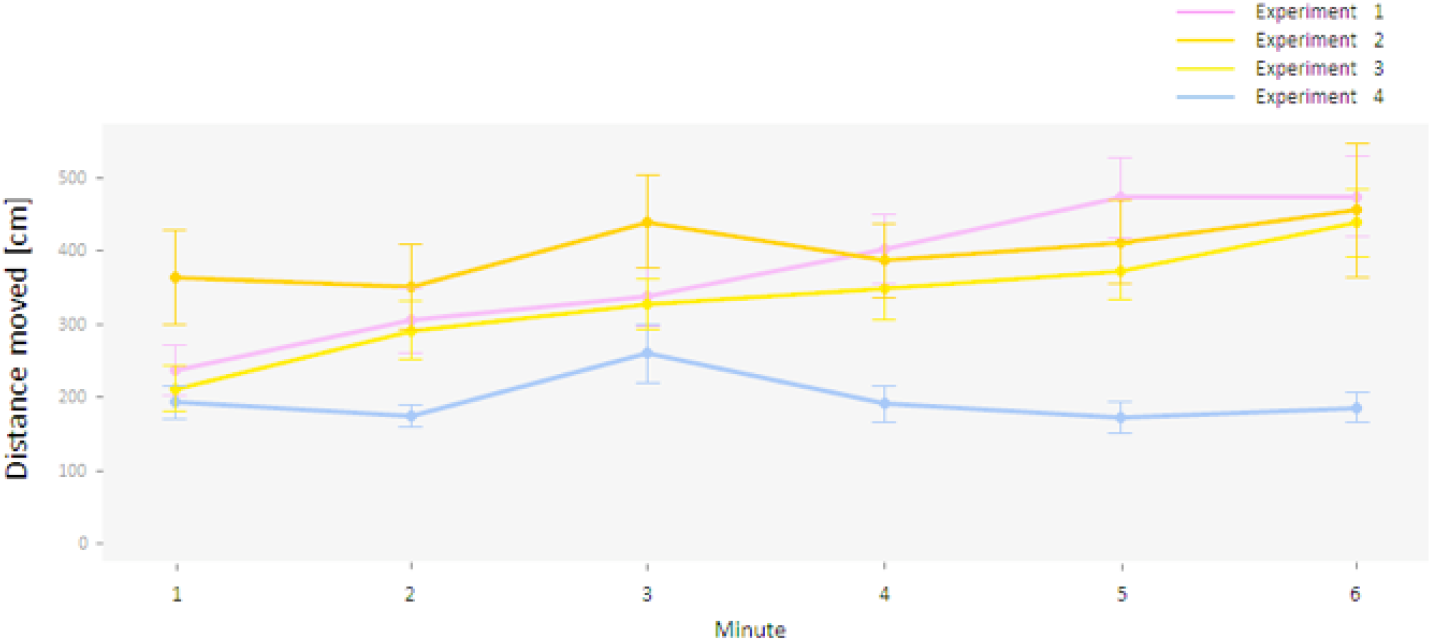
The lines show the distance moved by chicks in each experiment (mean ± SEM). Naïve chicks: Experiment 1; Imprinted chicks: same sex: Experiment 2; Imprinted chicks: opposite sex: Experiment 3; Imprinted chicks: mixed sex: Experiment 4

We compared the distance moved by chicks in the different experiments and found a significant effect of Experiment (F_3,476_ = 21.98; p<0.001). Post-hoc tests revealed a significant difference between Experiment 4 and Experiment 1 (Tukey’s test, p<0.001), Experiment 4 and Experiment 2 (Tukey’s test, p<0.001), and Experiment 4 and Experiment 3 (Tukey’s test, p<0.001). Fig. 4 shows that in Experiment 4 chicks were less active than in Experiment 1, 2 and 3.

**Figure 4.**
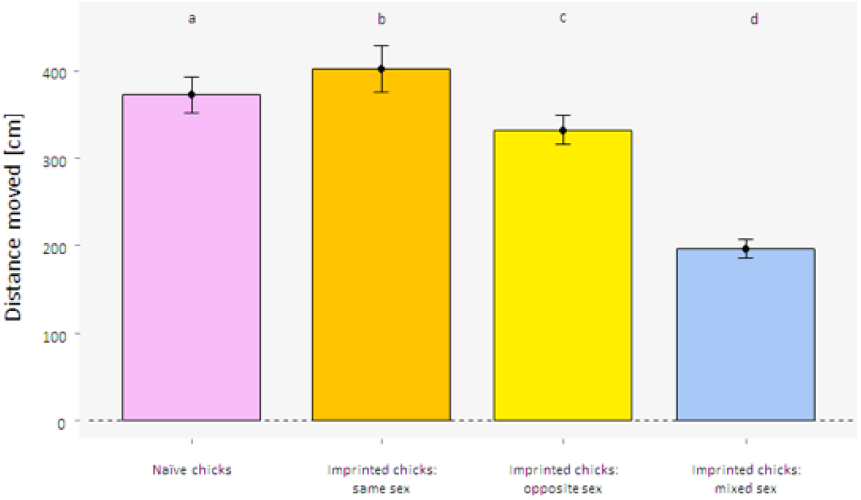
Boxplots showing the distance moved by chicks during the experiments (mean ± SEM). **a.** Naïve chicks: Experiment 1; **b.** Imprinted chicks: same sex: Experiment 2; **c.** Imprinted chicks: opposite sex: Experiment 3; ; **d.** Imprinted chicks: mixed sex: Experiment 4

### Body weight

Finally, when analysing the weight of males and females, we found no differences between sexes neither soon after hatching (t_123.62_ = −1.671, p = 0.097; mean for females 44.6 g, mean for males 46.0 g) nor after three days of imprinting (t_549.76_ = 0.856, p = 0.392; mean for females 70.6 g, mean for males 69.9 g).

## Discussion

We investigated whether domestic chicks (*Gallus gallus*) are able discriminate between sexes and prefer to aggregate with one sex or the other at hatching. To address this issue, we tested young chicks for their preference to approach male and female peers soon after birth or after three days of imprinting with peers of different sex. Our results show that soon after hatching (Experiment 1) chicks discriminate between sexes and preferentially approach female peers. This spontaneous preference is partially in line with the observations of Whiteside et al. (2017), who found that in the first week, female pheasants chicks preferred to associate with their own sex peers, while male pheasant chicks exhibited a random assortment.

At hatching, chicks appear sexually monomorphic, and in line with this we did not find significant differences in the weight of males and females. Analysing the distance moved in the experimental arena, we also observed no significant differences in the level of activity of the two sexes. These results suggest that visual features might have a limited role in sex discrimination at hatching. Although in our experiments we did not measure acoustic and olfactory variables, further experiments should investigate these modalities. Indeed, both acoustic (Gottlieb 1965; Park and Balaban 1991) and olfactory features (Caspers et al. 2017) drive parental attraction in absence of previous experience in other bird species, such as zebra finches (*Taeniopygia guttata,* Caspers et al. 2017), wood ducklings (*Aix sponsa,* Gottlieb 1965) and Japanese quails (*Coturnix coturnix japonica,* Park and Balaban 1991). In addition, although mainly observed in adult birds, sex recognition relies on both olfactory (e.g. in starlings, *Sturnus unicolor,* Amo et al. 2012 and crimson rosellas, *Platycercus elegans,* Mihailova et al. 2014) and acoustic (e.g. in Swinhoe’s Storm-Petrels, *Oceanodromam onorhis,* Taoka and Okumura 1990 and Eared Grebes, *Podiceps igricollis,* Nuechterlein & Buitron 1992) signals displayed by males and females. Further studies should clarify whether a similarity in the odour of female chicks and the mother hen underlie chicks’ attraction for female chicks in absence of the mother.

When we analysed the preference of active chicks during the six minutes of Experiment 1, we found that male and female chicks modified their preference for females with time, with a peak in the fourth minute. Changes in the preference show the tendency of chicks to explore more than one stimulus when they are exposed for the first time to chicks of different sex. This changing preference has also been observed by Versace et al. (2017a). Here authors found that different breeds differ in their propensity to aggregate with novel stimuli in the subsequent 5 minutes after the exposure, with Polverara breed maintaining a constant preference throughout the test and Padovana and Robusta breed progressively exploring the novel stimuli. Similar exploratory behaviour for different stimuli has been detected during the imprinting process and has been interpreted as a part of the formation of parental representation (Bateson 1973; Vallortigara and Andrew 1991, 1994; Jackson and Bateson 1974). The observe changes in preference may therefore be an early strategy connected to the imprinting process.

After documenting the precocious attraction for females in newly hatched chicks, we tested the role of experience on sex preference during chicks’ development. We thus manipulated the rearing conditions of the animals by imprinting the chicks with peers of different sex combinations. We observed that three-day old chicks reared with four same-sex chicks (Experiment 2), four opposite-sex chicks (Experiment 3) or two same-sex and two opposite-sex (Experiment 4) chicks did not show any preference to aggregate with males or females. These findings suggest that the preference for females observed at birth is not stable but limited to the first stages soon after hatching. An exception was found for active chicks imprinted with same-sex chicks (Experiment 2). Compared to Experiment 3 and 4, the preference towards females in active chicks of Experiment 2 did not completely disappear, given that males in the first minute still preferred to aggregate with females. The persistence of this predisposition may find its roots in the “slight novelty effect” (Jackson and Bateson 1974; Bateson 1979; Versace et al. 2006) and is in line with previous studies observing a preference of males towards novel stimuli (Vallortigara and Andrew 1994). Similarly, in Experiment 2 females represented a novel stimulus for males, who have been previously exposed to same sex chicks only. Accordingly, the preference for females disappeared when chicks were reared with opposite sex chicks (Experiment 3) and mixed sex chicks (Experiment 4).

The decrease of this spontaneous preference after three days of imprinting suggests the existence of a transient time window for its expression, as it has been observed for other social predispositions, such as the early preference for objects that move changing speed (Versace et al. 2019) and the preference to approach a stuffed-hen compared to a scrambled hen, that is stronger in 24 hour-old than in two-day old chicks (Johnson et al. 1985; Johnson and Horn 1988). Mounting evidence supports the hypothesis that early predispositions are not constant through the life course. This changing temporal course of early predispositions is not restricted to the fowl (see Rosa-Salva et al. under review for a thorough discussion on the adaptive role of the temporal dynamics in early predispositions). In human infants, for instance, a fluctuation in the response to stimuli of different nature, such as sound sources (Field et al. 1980; Muir et al. 1989) and face-like patterns (Johnson et al. 1991; Simion 1998), has been observed during the development.

A further variable that could influence the development of sex preference is the number of peers used during the

The exploration of the cognitive processes underpinning the emergence of sexual preferences is essential to understand social behaviour, particularly in sexually dimorphic animals. Overall, our study shows that inexperienced chicks are predisposed to aggregate with females at hatching, and that this preference quickly fades during the development, independently from the exposure to chicks of different sex. Further studies should explore the mechanisms at the basis of sex discrimination, and the changes that determine a transient window in the expression of early predispositions.

imprinting phase, although in previous studies its effect has been detected after several days of imprinting and tested in relation to the mating period (Adkins-Regan and Krakauer 2000; Leonard et al.1993; Madden and Whiteside 2013). The variation in sex ratio during the rearing phase has also been shown to have an effect on the ability to discriminate between sexes, affecting differently males and females in their preference for mating partners, with females needing a higher number of opposite sex peers to discriminate males from females (Bateson 1980). Therefore, we cannot exclude that the number of males and females in the imprinting conditions of Experiments 2, 3 and 4 were not sufficient to modulate the initial social preference shown by chicks. To further explore the effect of the sex ratio of imprinting stimuli on social predispositions further experiments should test chicks after different duration of exposure to peers as well as to vary the proportion of males and females used as imprinting stimuli.

The findings of our imprinting experiments (Experiments 2-4) suggest a lack of sexual preference in chicks of both sexes when their peers are not sexually dimorphic, as confirmed by the similarity of three-days-old males and females in their body weight and physical activity. We thus suggest that sexual segregation has a limited role in driving chicks aggregation in the first week of life, except for the first hours immediately after hatching, before filial imprinting takes place. Such an absence of sexual assortment in the first days of life may be important to enhance the social cohesion of the flock as antipredator response, postponing the sexual segregation to a later emergence of physical differences between sexes. In line with this idea, in pheasants of both sexes the increased predisposition to aggregate with same sex peers corresponded to an increase in sexual dimorphism of chicks, highlighting the importance of physical features in driving sexual segregation later in life (Whiteside et al. 2017).

We did not find differences between males and females in the distance moved but we observed that chicks were less active in Experiment 4 compared to Experiment 1, 2 and 3. The reduced distance moved in Experiment 4 may be due to the rearing conditions of chicks before the testing phase, where chicks were exposed for three days to both males and females. Therefore, a low motivation to explore the novel stimuli during the testing phase could be explained as a lack of novelty effect related to the sex of the stimuli.

## Compliance with ethical standards

### Conflict of interest

The authors declare that they have no conflict of interest.

### Ethical approval

All experiments comply with the ASAB/ABS Guidelines for the Use of Animals in Research and with the current Italian and European Community laws for the ethical treatment of animals and the experimental procedures were approved by the Ethical Committee of University of Trento and licensed by the Italian Health Ministry (permit number 630-2018 PR).

## Supplementary materials

**Supplementary Figure 1.**
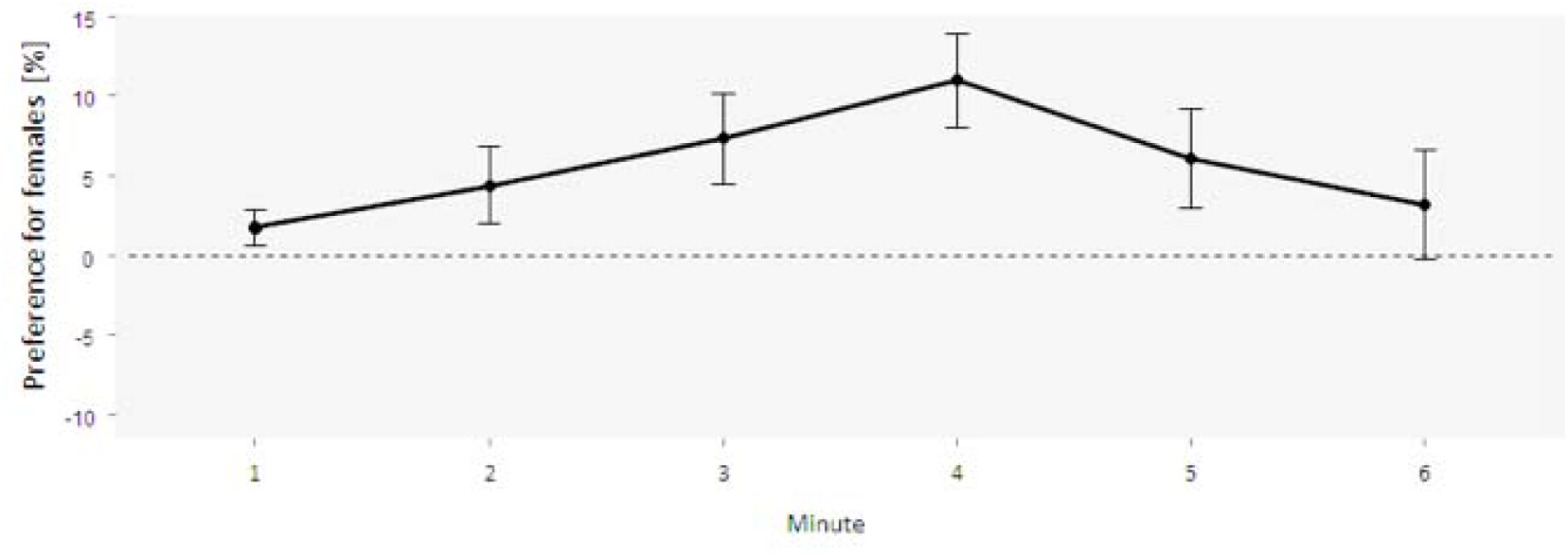
The line shows the preference for females of naïve chicks in Experiment 1, across the six minutes of test (mean ± SEM)

**Supplementary Figure 2.**
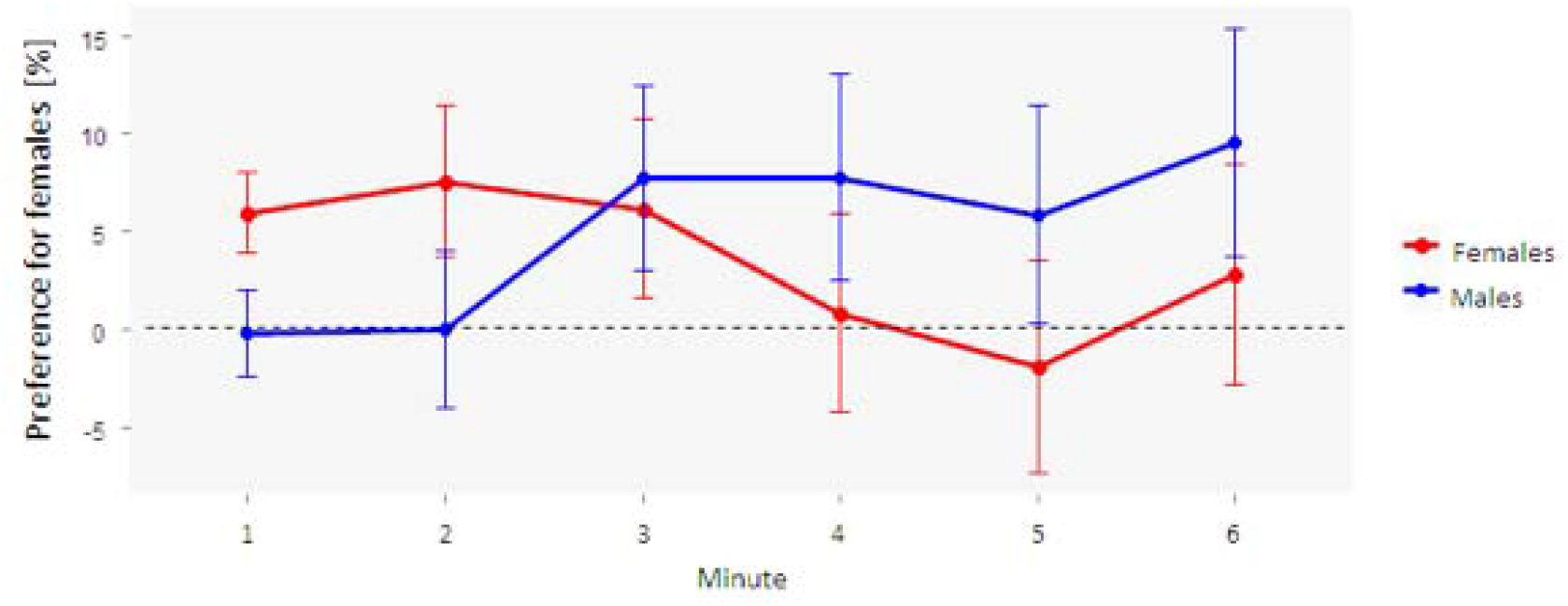
The lines show the preference for females exhibited by male and female chicks imprinted with same-sex peers in Experiment 2, across the six minutes of test (mean ± SEM)

**Supplementary Figure 3.**
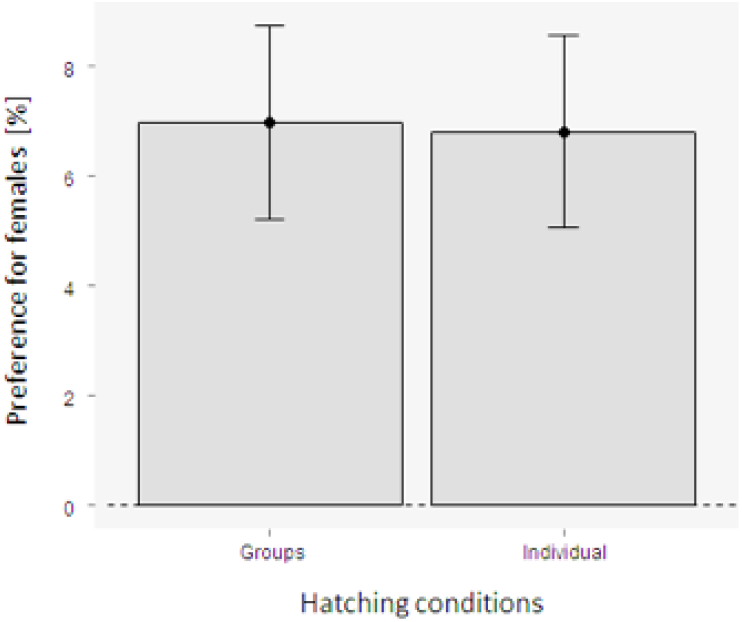
Bar plots showing the preference for females of chicks hatched in individual plastic box or in groups (mean ± SEM). We did not find any differences between the conditions in which chicks hatched in individual boxes (11 x 8.5 x 14 cm) and chicks hatched in groups (t = 0.388, df = 128.4, p = 0.969).

## Notes

### Competing Interest Statement

The authors have declared no competing interest.

## References

Adkins-Regan, E, Krakauer, A. (2000). Removal of adult males from the rearing environment increases preference for same-sex partners in the zebra finch. Anim Behav 60: 47–53. https://doi.org/10.1006/anbe.2000.1448

Amo L, Avilés JM, Parejo D, Peña A, Rodríguez J, Tomás G (2012) Sex recognition by odour and variation in the uropygial gland secretion in starlings. J Anim Ecol 81: 605–613. https://doi.org/10.1111/j.1365-2656.2011.01940.x

Bateson PPG (1966) The characteristics and context of imprinting. Biol Rev 41: 177–217.

Bateson PPG (1973) Preferences for familiarity and novelty: a model for the simultaneous development of both. J Theor Biol 41: 249–259. https://doi.org/10.1016/0022-5193(73)90117-3

Bateson PPG (1979) How do sensitive periods arise and what are they for?. Anim Behav 27: 470–486. https://doi.org/10.1016/0003-3472(79)90184-2

Bateson P (1980) Optimal outbreeding and the development of sexual preferences in Japanese quail. Z Tierpsychol 53: 231–244. https://doi.org/10.1111/j.1439-0310.1980.tb01052.x

Bateson P (1990) Is imprinting such a special case?. Philos T Roy Soc B 329: 125–131. https://doi.org/10.1098/rstb.1990.0157

Bischof HJ (2003) Neural mechanisms of sexual imprinting. Anim Biol 53: 89–112. https://doi.org/10.1163/157075603769700313

Bolhuis JJ (1991) Mechanisms of avian imprinting: a review. Biol Rev 66: 303–345. https://doi.org/10.1111/j.1469-185X.1991.tb01145.x

Caspers BA, Hagelin JC, Paul M, Bock S, Willeke S, Krause ET (2017) Zebra Finch chicks recognize parental scent, and retain chemosensory knowledge of their genetic mother, even after egg cross-fostering. Sci Rep-Uk 7: 18. https://doi.org/10.1038/s41598-017-13110-y.

Cheng KM, Shoffner RN, Phillips RE, Lee FB (1979) Mate preference in wild and domesticated (game-farm) mallards: II. Pairing success. Anim Behav 27: 417–425. https://doi.org/10.1016/0003-3472(79)90177-5

Clayton NS (1989) Song, sex and sensitive phases in the behavioural development of birds. Trends Ecol Evol 4: 82–84. https://doi.org/10.1016/0169-5347(89)90156-0

Di Giorgio E, Loveland JL, Mayer U, Rosa-Salva O, Versace E, Vallortigara G (2017) Filial responses as predisposed and learned preferences: Early attachment in chicks and babies. Behav Brain Res 325: 90–104. https://doi.org/10.1016/j.bbr.2016.09.018

Field J, Muir DW, Pilon R, Sinclair M, Dodwell PC (1980) Infants’ orientation to lateral sounds from birth to three months. Child Dev 51: 295–298. DOI: 10.2307/1129628

Gottlieb G (1965) Imprinting in relation to parental and species identification by avian neonates. J Comp Physiol Psych 59: 345. https://doi.org/10.1037/h0022045

Gottlieb G (1974) On the acoustic basis of species identification in wood ducklings (Aix sponsa). J Comp Physiol Psych 87: 1038–1048. https://doi.org/10.1037/h0037601

Hess EH (1956) Natural preferences of chicks and ducklings for objects of different colors. Psychol Rep 2: 477–483. doi:10.2466/pr0.2.7.477-48

Hess EH, Gogel WC (1954) Natural preferences of the chick for objects of different colors. J Psychol 38: 483–493. doi:10.1080/00223980.1954.9712955

Horn G (2004) Pathways of the past: the imprint of memory. Nat Rev Neurosci 5: 108–120. http://dx.doi.org/10.1038/nrn1324.

Jackson PS, Bateson, PPG (1974) Imprinting and Exploration of Slight Novelty in Chicks. Nature 251: 609–610. https://doi.org/10.1038/251609a0

Johnson MH, Bolhuis JJ, Horn G (1985) Interaction between acquired preferences and developing predispositions during imprinting. Anim Behav 33: 1000–1006. https://doi.org/10.1016/S0003-3472(85)80034-8

Johnson MH, Horn G (1988) Development of filial preferences in dark-reared chicks. Anim Behav 36: 675–683. https://doi.org/10.1016/S0003-3472(88)80150-7

Johnson MH, Dziurawiec S, Ellis H, Morton J (1991) Newborns’ preferential tracking of face-like stimuli and its subsequent decline. Cognition 40: 1–19. https://doi.org/10.1016/0010-0277(91)90045-6

Launay F, Mills AD, Faure JM (1993) Effects of test age, line and sex on tonic immobility responses and social reinstatement behaviour in Japanese quail *Coturnix japonica*. Behav Process 29: 1–16.https://doi.org/10.1016/0376-6357(93)90023-K

Leonard ML, Zanette L, Thompson BK, Fairfull RW (1993) Early exposure to the opposite sex affects mating behaviour in White Leghorn chickens. Appl Anim Behav Sci 37: 57–67. https://doi.org/10.1016/0168-1591(93)90070-6

Madden JR, Whiteside MA (2013) Variation in female mate choice and mating success is affected by sex ratio experienced during early life. Anim Behav 86: 139–142. https://doi.org/10.1016/j.anbehav.2013.05.003

Matsushima T, Izawa EI, Aoki N, Yanagihara S (2003) The mind through chick eyes: memory, cognition and anticipation. Zool Sci 20: 395–408. http://dx.doi.org/10.2108/zsj.20.395.

McCabe BJ (2013) Imprinting. Wires Cogn Sci 4: 375–390. http://dx.doi.org/10.1002/wcs.1231.

Mihailova M, Berg ML, Buchanan KL, Bennett AT (2014) Odour-based discrimination of subspecies, species and sexes in an avian species complex, the crimson rosella. Anim Behav 95: 155–164.https://doi.org/10.1016/j.anbehav.2014.07.012

Miura M, Nishi D, Matsushima T (2020) Combined predisposed preferences for colour and biological motion make robust development of social attachment through imprinting. Anim Cogn 23: 169–188. https://doi.org/10.1007/s10071-019-01327-5

Muir DW, Clifton RK, Clarkson MG (1989) The development of a human auditory localization response: A U-shaped function. Can J Psychology 43: 199–216. https://doi.org/10.1037/h0084220

Nuechterlein GL, Buitron D (1992) Vocal advertising and sex recognition in eared grebes. Condor 94: 937–943. https://doi.org/10.2307/136929

Park T, Balaban E (1991) Relative salience of species maternal calls in neonatal gallinaceous birds: a direct comparison of Japanese quail *(Coturnix coturnix japonica)* and domestic chickens *(Gallus gallus domesticus)*. J Comp Psychol 105: 45. https://doi.org/10.1037/0735-7036.105.1.45

Rosa-Salva O, Regolin L, Vallortigara G (2010) Faces are special for newly hatched chicks: evidence for inborn domain-specific mechanisms underlying spontaneous preferences for face-like stimuli. Developmental Sci 13: 565–577. https://doi.org/10.1111/j.1467-7687.2009.00914.x

Rosa-Salva O, Grassi M, Lorenzi E, Regolin L, Vallortigara G (2016) Spontaneous preference for visual cues of animacy in naïve domestic chicks: The case of speed changes. Cognition 157: 49–60. https://doi.org/10.1016/j.cognition.2016.08.014

Rosa-Salva O, Mayer U, Vallortigara G (2019) Unlearned visual preferences for the head region in domestic chicks. PLoS One 14: e0222079. https://doi.org/10.1371/journal.pone.0222079

Rosa-Salva O, Mayer U, Versace E, Hebert M, Lemaire BS, Vallortigara G (Under review) Sensitive periods for social development: Interactions between predisposed and learned mechanisms.

Sherrod L (1974) The role of sibling associations in the formation of social and sexual companion preferences in ducks *(Anas platyrhynchos):* An investigation of the ‘Primacy versus Recency’ question. Z Tierpsychol 34: 247–264. https://doi.org/10.1111/j.1439-0310.1974.tb01801.x

Simion F, Valenza E, Umilta C, Barba BD (1998) Preferential orienting to faces in newborns: A temporal–nasal asymmetry. J Exp Psychol Human 24: 1399–1405. https://doi.org/10.1037/0096-1523.24.5.1399

Taoka M, Okumura H (1990) Sexual differences in flight calls and the cue for vocal sex recognition of Swinhoe’s Storm-Petrels. Condor 92: 571–575. https://doi.org/10.2307/1368674

ten Cate C (1986) Does behavior contingent stimulus movement enhance filial imprinting in Japanese quail?. Dev Psychobiol 19: 607–614. https://doi.org/10.1002/dev.420190611

Vallortigara G (1992) Affiliation and aggression as related to gender in domestic chicks (Gallus gallus). J Comp Psychol 106: 53–57. https://doi.org/10.1037/0735-7036.106.1.53

Vallortigara G, Andrew RJ (1991) Lateralization of response by chicks to change in a model partner. Anim Behav 41: 187–194. https://doi.org/10.1016/S0003-3472(05)80470-1

Vallortigara G, Andrew RJ (1994) Differential involvement of right and left hemisphere in individual recognition in the domestic chick. Behav Process 33: 41–57. doi:10.1016/0376-6357(94)90059-0

Vallortigara G, Regolin L (2006) Gravity bias in the interpretation of biological motion by inexperienced chicks. Curr Biol 16: 279–280. doi:10.1016/j.cub.2006.03.052

Vallortigara G, Versace E (2018) Filial Imprinting. Encyclopedia of Animal Behavior, 1943–1948.

Vallortigara G, Cailotto M, Zanforlin M (1990) Sex differences in social reinstatement motivation of the domestic chick *(Gallus gallus)* revealed by runway tests with social and nonsocial reinforcement. J Comp Psychol 104: 361–367. https://doi.org/10.1037/0735-7036.104.4.361

Vallortigara G, Regolin L, Marconato F (2005) Visually inexperienced chicks exhibit spontaneous preference for biological motion patterns. PLoS Biol 3: e208. doi: 10.1371/journal.pbio.0030208

Versace E, Regolin L, Vallortigara G (2006) Emergence of grammar as revealed by visual imprinting in newly-hatched chicks. The evolution of language. In: Proceedings of the 6th international conference, Rome, 12–15 April 2006.

Versace E, Fracasso I, Baldan G, Dalle Zotte A, Vallortigara G (2017a) Newborn chicks show inherited variability in early social predispositions for hen-like stimuli. Sci Rep-Uk 7: 40296. https://doi.org/10.1038/srep40296

Versace E, Spierings MJ, Caffini M, Ten Cate C, Vallortigara G (2017b) Spontaneous generalization of abstract multimodal patterns in young domestic chicks. Anim Cogn 20: 521–529. https://doi.org/DOI10.1007/s10071-017-1079-5

Versace E, Ragusa M, Vallortigara G (2019) A transient time window for early predispositions in newborn chicks. Sci Rep-Uk 9: 1–7. https://doi.org/10.1038/s41598-019-55255-y

Whiteside MA, van Horik JO, Langley EJ, Beardsworth CE, Laker PR, Madden JR (2017) Differences in social preference between the sexes during ontogeny drive segregation in a precocial species. Behav Ecol Sociobiol 71: 103. https://doi.org/10.1007/s00265-017-2332-2

Whiteside MA, van Horik JO, Langley EJ, Beardsworth CE, Madden JR (2018) Size dimorphism and sexual segregation in pheasants: tests of three competing hypotheses. PeerJ 6: e5674. DOI 10.7717/peerj.5674

Whiteside MA, van Horik JO, Langley EJ, Beardsworth CE, Capstick LA, Madden JR (2019) Patterns of association at feeder stations for Common Pheasants released into the wild: sexual segregation by space and time. Ibis 161: 325–336. https://doi.org/10.1111/ibi.12632

Worsley, A. (1974). Long-term Effects of Imprinting Exposure upon Breed Discriminatory Behaviour in Chickens: I. Imprinting to peers (“peerprinting”). Zeitschrift für Tierpsychologie, 35(1), 1–9.

